# SparkINFERNO: A scalable high-throughput pipeline for inferring molecular mechanisms of non-coding genetic variants

**DOI:** 10.1101/2020.01.07.897579

**Authors:** Pavel P. Kuksa, Chien-Yueh Lee, Alexandre Amlie-Wolf, Prabhakaran Gangadharan, Elizabeth E. Mlynarski, Yi-Fan Chou, Han-Jen Lin, Heather Issen, Emily Greenfest-Allen, Otto Valladares, Yuk Yee Leung, Li-San Wang

## Abstract

**Summary:** We report SparkINFERNO (Spark-based INFERence of the molecular mechanisms of NOn-coding genetic variants), a scalable bioinformatics pipeline characterizing noncoding GWAS association findings. SparkINFERNO prioritizes causal variants underlying GWAS association signals and reports relevant regulatory elements, tissue contexts, and plausible target genes they affect. To achieve this, the SparkINFERNO algorithm integrates GWAS summary statistics with large-scale collection of functional genomics datasets spanning enhancer activity, transcription factor binding, expression quantitative trait loci, and other functional datasets across more than 400 tissues and cell types. Scalability is achieved by an underlying API implemented using Apache Spark and Giggle-based genomic indexing. We evaluated SparkINFERNO on large GWAS studies and show that SparkINFERNO is more than 60-times efficient and scales with data size and amount of computational resources.

**Availability:** SparkINFERNO runs on clusters or a single server with Apache Spark environment, and is available at https://bitbucket.org/wanglab-upenn/SparkINFERNO or https://hub.docker.com/r/wanglab/spark-inferno.

## 1 Introduction

Genome-wide association studies (GWASs) have successfully identified over 70,000 genetic variants associated with more than 3,000 human diseases and phenotypes (Buniello *et al.*, 2019). Interpretation of these associations remain difficult (Watanabe *et al.*, 2017; Amlie-Wolf *et al.*, 2018) as most GWAS hits are in the noncoding genome. Resolution of GWAS is limited as neighboring variants have similar associations due to linkage disequilibrium (LD) (Amlie-Wolf *et al.*, 2018). Our recently developed INFERNO method (Amlie-Wolf *et al.*, 2018) focuses on identifying potentially causal variants underlying observed GWAS associations by integrating with hundreds of functional genomics datasets. The current INFERNO implementation is not optimized for big data, and a scalable framework for annotating genetic variants and genomic regions generated by various human genetic studies in a high throughput manner is in need for systematic large scale genomic and genetic analyses.

The scale and heterogeneity of functional genomics datasets and annotations necessitate systematic, integrative analysis and interpretation of GWAS association findings. For example, while INFERNO (Amlie-Wolf *et al.*, 2018) uses relatively small set of functional genomics datasets, projects such as GTEx (Aguet *et al.*, 2017), FANTOM5 (Andersson *et al.*, 2014), ENCODE (Bernstein *et al.*, 2012) and Roadmap Epigenomics (Consortium *et al.*, 2015) produce >60,000 experimental datasets across >1,100 tissues, cell types, biological conditions, each with millions to billions of records across the genome. In order to pair these functional annotations with modern population-level studies such as UK Biobank (500,000 individuals with >2,500 phenotypes), we need a scalable, high-throughput, robust and easy to use software that can systematically interpret hundreds of millions of genotypes across millions of participants.

We implemented SparkINFERNO as a scalable, high-throughput automated workflow that integrates a large-scale functional genomics data repository and processes GWAS results by performing LD analysis, functional evidence evaluation and aggregation, Bayesian co-localization analysis of GWAS and eQTL signals, characterize the downstream regulatory effects including the tissue contexts, regulatory elements, and target genes that they affect. We applied SparkINFERNO on inflammatory bowel disease (IBD) (Liu *et al.*, 2015) and the International Genomics of Alzheimer’s disease (AD) GWAS datasets (Lambert *et al.*, 2013) and show that this scalable framework is at least 60-times more efficient and able to identify the molecular mechanisms underlying noncoding GWAS signals.

## 2 Methods

We chose Apache Spark (Zaharia *et al.*, 2016) and Python for a scalable implementation of INFERNO (Amlie-Wolf *et al.*, 2018) (see Supplementary Table 1). The new SparkINFERNO is highly scalable, modular, and coupled with an integrated functional genomics data repository (Figure 1; Supplementary Figure S1; Supplementary Tables S1, S2). Analysis modules perform various types of genomic data integration to produce functional evidence including tissue-specific regulatory elements (enhancers), transcription factor (TF) activity, chromatin states, and genetic regulation (eQTL) information. SparkINFERNO implements scalable genomic querying (Supplementary Figures S2, S3) using Spark parallel transformations and Giggle-based genomic indexing (Layer *et al.*, 2018). SparkINFERNO can be extended with additional annotation data and/or customized evaluation modules. Results are reported by individual evaluation modules and as combined summaries (Supplementary Methods).

**Fig. 1.**
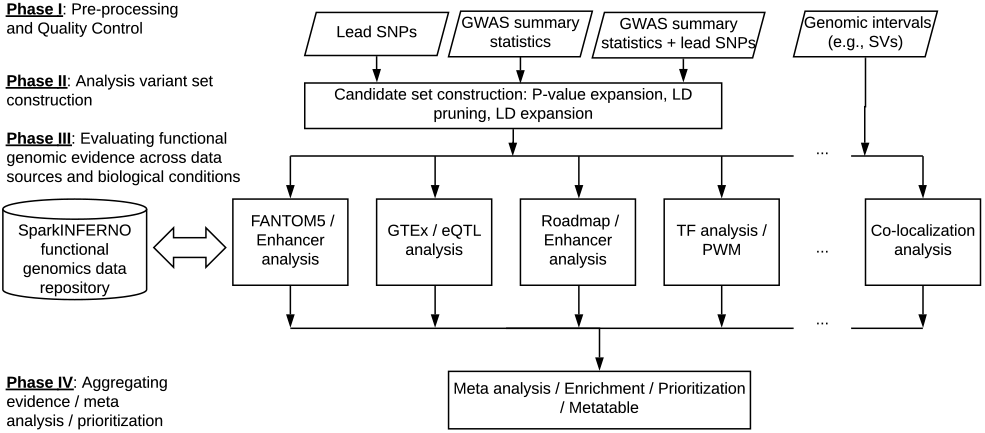
Overview of SparkINFERNO

SparkINFERNO accepts complete GWAS summary statistics or top GWAS association variants as the input and generates a list of potentially causal variants, affected tissue-specific enhancers and target gene(s) as the output. The entire workflow consists of four phases (Figure 1): (1) **Preprocessing** and QC of GWAS input; (2) **Generating candidate set** of potentially causal variants, (3) **Evaluating functional genomic evidence** across genomic datasets in a tissue-specific manner including regulatory elements (enhancers); eQTL co-localization, transcriptional factor binding sites (TFBSs) and others for each GWAS locus/signal; (4) **Aggregating evidence** to infer prioritization of causal variants, including information on affected tissues/cell types, regulatory elements, TFs, and target genes. See Supplementary Methods for technical details.

The Pre-Processing phase takes raw GWAS summary statistics in a TSV format as input, resolves reference and alternative alleles, check allele frequencies in the reference population (e.g., 1000 Genomes Project), and produce quality control flags. Quality control steps mark GWAS variants with inconsistent alleles that could not be matched with reference genotype data (Supplementary Figure S4 and Supplementary Methods).

The Candidate Set Construction phase expands genome-wide significant associations into a putative causal variant set by pruning significant variants into a smaller set of independent variants using publicly available LD data (e.g. 1000 Genome), and then expanding these signals into putative causal sets consisting of nearby variants in LD. The user can specify the reference population in LD pruning/expansion to match the population underlying the input GWAS study. Supplementary Methods and Figure S4 provides details of the workflow for generating putative variant sets.

The Evaluation phase executes Spark-based annotation jobs in parallel (Figure 1). SparkINFERNO uses an integrated repository of annotations for genomic elements (promoters, exons, introns, etc.), non-coding RNAs, regulatory elements such as enhancers, TFBSs, and others (integrated data and data repository implementation in Supplementary Table S2 and Figure S1). The current SparkINFERNO implementation contains 3.5 billion genomic intervals from 2,342 tracks for 32 tissue categories.

In the final Aggregation phase, SparkINFERNO combines functional evidence from individual genomic analyses and produces a list of candidate variants, enhancer elements, and genes with which they interact as supported by FANTOM5, Roadmap, GTEx, TF binding and other functional evidence. SparkINFERNO performs co-localization analysis (Supplementary Figure S5) of the GWAS and eQTL signals across genome-wide significant loci using COLOC (Giambartolomei *et al.*, 2014).

To install SparkINFERNO, users can either install the package (https://bitbucket.org/wanglab-upenn/SparkINFERNO) on their own Spark cluster, or use a pre-created Docker image (wanglab/spark-inferno). To run SparkINFERNO, the user first edits the configuration file and provides input GWAS specifications. A complete run of SparkINFERNO produces candidate potentially causal variants, target genes, tissue contexts, regulatory elements, and detailed BED files documenting overlaps with functional genomics and annotation datasets.

## 3 Results

We evaluated SparkINFERNO on our AWS Spark cluster using publicly available IBD and IGAP AD GWAS datasets containing 11,555,676 and 8,080,502 variants respectively. For the IGAP GWAS dataset, SparkINFERNO took 993 seconds on a 16-core Linux server to complete the analysis, whereas the original INFERNO took 60,973 seconds. SparkINFERNO is 61-times faster (Supplementary Figure S2) and scales well with the amount of computational resources (Supplementary Figure S3). SparkINFERNO identified 1,418 and 15,343 candidate causal variants and 149 and 1,002 co-localized target gene-tissue combinations for IGAP and IBD, respectively. As can be seen from distribution of identified overlaps across functional genomics datasets and tissue types (Supplementary Figures S6 and S7) SparkINFERNO identifies genes and tissues that are likely important for the disease etiology.

