## Supplementary Information for "SparkINFERNO: A scalable high-throughput pipeline for inferring molecular mechanisms of non-coding genetic variants"

### Supplementary Tables

**Table S1.** Improvements provided by SparkINFERNO

| **Features** | **SparkINFERNO** | **INFERNO** |
|---|---|---|
| Distributed, parallel implementation (Apache Spark) | ✓ | - |
| Scalable genomic data search engine (Spark+Giggle) | ✓ | - |
| Integrated functional genomics data collection: indexed, annotated, searchable, extensible; with scalable high-throughput interface | ✓ | - |
| Scalable overlap with (thousands of) genome-wide annotations | ✓ | - |
| Scalable co-localization analysis | ✓ | - |
| Scalable enrichment analysis:<br>per-track/dataset, per-tissue/cell type, per-annotation category | ✓ | - |
| Scalable transcription factor binding (TFBS) analysis | ✓ | - |
| Modular, extensible architecture of evidence aggregating meta-table | ✓ | |
| Support for local, cluster, and cloud computing platforms | ✓ | Local/bsub only |
| Support for both GRCh38/hg38 and GRCh37/hg19 genome builds | ✓ | - |
| Modular, extensible, parallel architecture; support for custom analysis modules | ✓ | - |
| GWAS pre-processing and quality control (QC) | ✓ | - |

**Table S2.** List of functional genomic datasets integrated into SparkINFERNO data repository. All of integrated datasets are categorized into a common set of broader tissue and cell categories (Amlie-Wolf *et al.*, 2018) for integration with SparkINFERNO

| Data source | Description | #tracks | #genomic features |
|---|---|---|---|
| **Data for GRCh37/hg19 genome build** | | | |
| DASHR2 | Small RNA loci | 632 | 5,773,364 |
| DASHR2_small_RNA_Genes | Small RNA gene annotations | 2 | 1,378,260 |
| FANTOM5 | Enhancer loci | 112 | 197,373 |
| GTEx_v6p_all_association | eQTL (all associations) | 44 | 419,398,672 |
| GTEx_v6p_signif_association | eQTL (significant associations) | 44 | 26,393,285 |
| GTEx_v7_all_association | eQTL (all associations) | 48 | 36,781,356 |
| GTEx_v7_signif_association | eQTL (significant associations) | 48 | 588,260,861 |
| Homer | Transcription factor binding sites (TFBS) | 1 | 568,191,857 |
| Inferno_genomic_partition | Genomic partion (mRNA, lncRNA, other genes, repeats) | 3 | 6,030,654 |
| ROADMAP_Enhancers | Enhancers (ChromHMM) | 127 | 13,617,978 |
| TargetScan_v7p2 | miRNA target | 9 | 12,467,265 |
| **Data for GRCh38/hg38 genome build** | | | |
| DASHR2 | Small RNA loci | 632 | 5,715,577 |
| DASHR2_small_RNA_Genes | Small RNA gene annotations | 2 | 1,529,342 |
| GTEx_v8_all_association | eQTL (all associations) | 48 | 694,223,963 |
| GTEx_v8_signif_association | eQTL (significant associations) | 48 | 71,478,479 |
| Homer | TFBS | 1 | 595,761,650 |
| Inferno_genomic_partition | Genomic partion (mRNA, lncRNA, other genes, repeats) | 3 | 6,247,462 |
| ROADMAP_Enhancers | Enhancers (ChromHMM) | 127 | 13,613,105 |
| FANTOM5-lifted* | Enhancer loci | 112 | 197,287 |
| TargetScan_v7p2-lifted* | miRNA target | 9 | 12,465,465 |
| | **Total** | **2,342** | **3,515,846,585** |

* These data sources were lifted from hg19 to hg38 reference. Liftover was performed on all of the data sources for which no hg38 data was available. Hg19 genome coordinates are lifted using UCSC liftOver utility hg19ToHg38.over.chain.gz. Hg19 coordinates are retained in the lifted-over files.

### Supplementary Figures

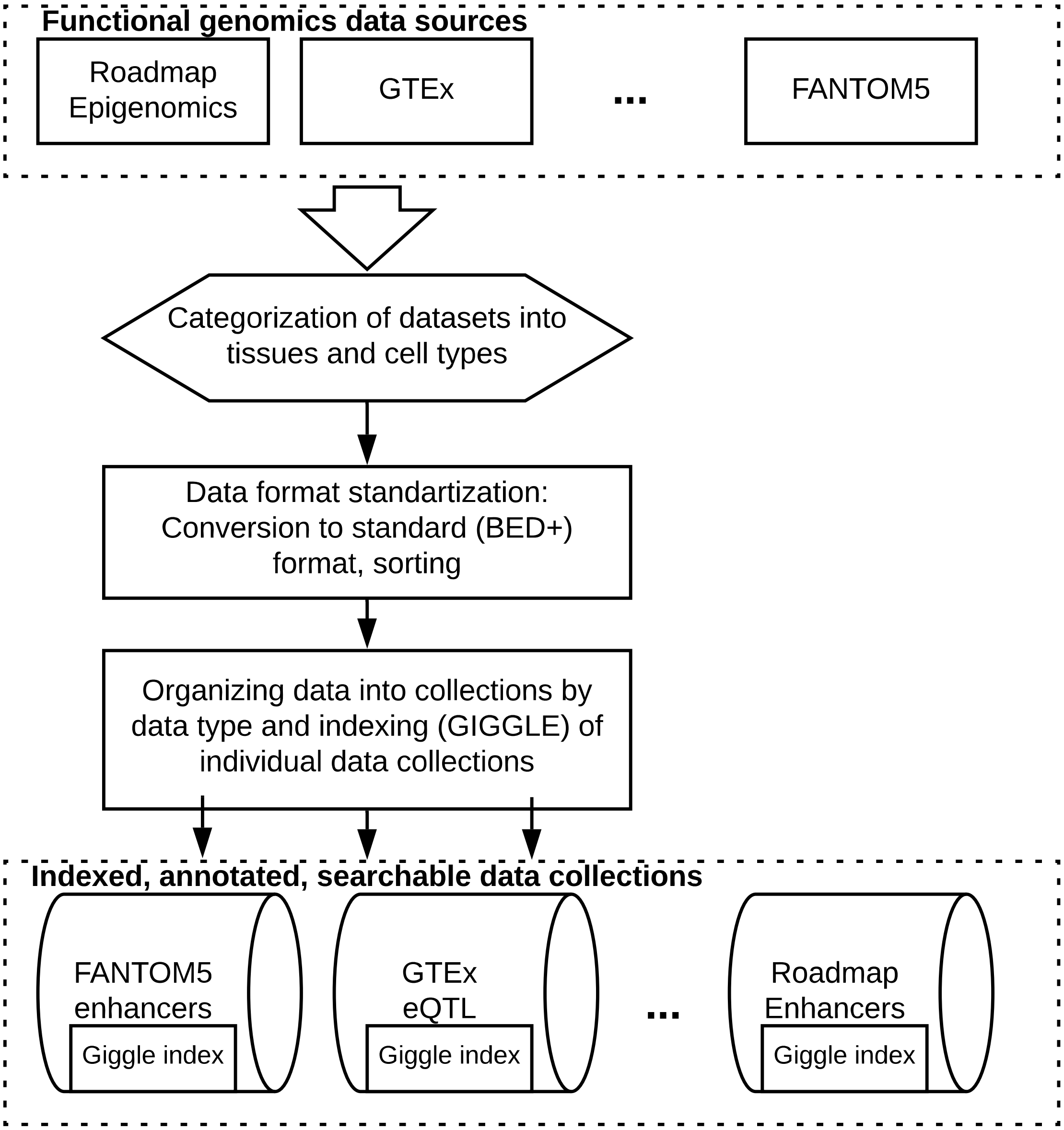


**Figure S1.** SparkINFERNO data repository architecture. All Integrated functional genomics datasets were grouped into broader cell and tissue categories using OBO Foundry ontologies. All these genomic tracks were transformed into BED format and organized into data collections by data type and data source. Each data collection was indexed using GIGGLE (https://github.com/ryanlayer/giggle). All genomic tracks are available for both GRCh37/hg19 and GRCh38/hg38 reference genomes.

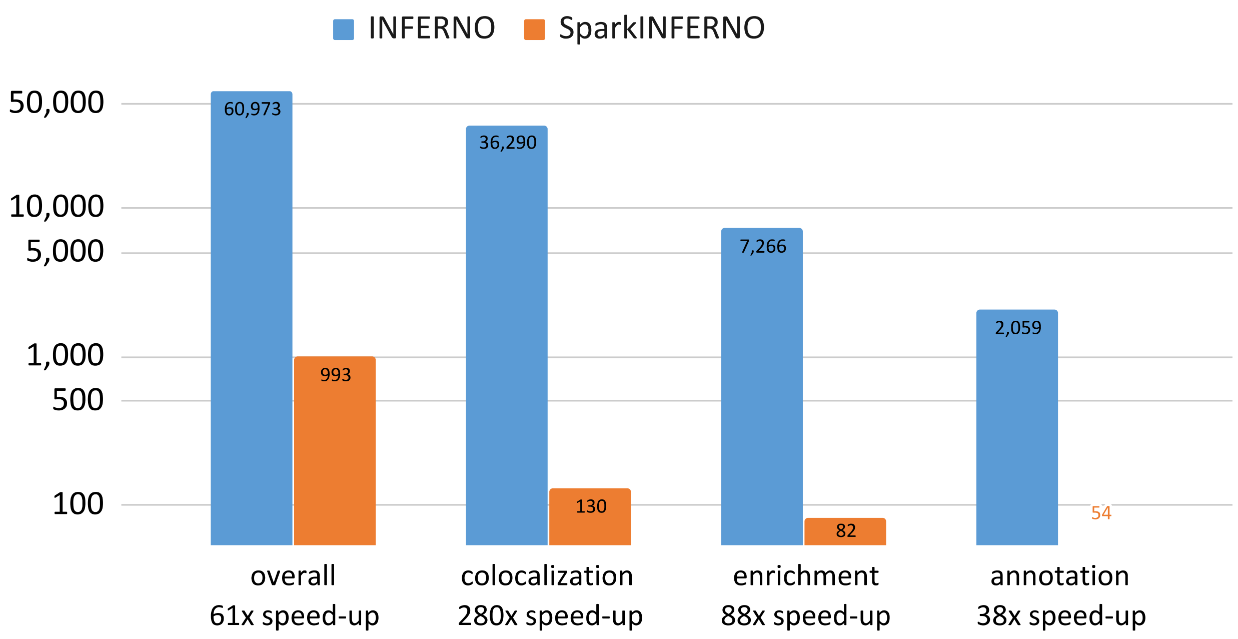


**Figure S2.** Running time comparison for Spark-INFERNO and INFERNO. Note the logarithmic scale of Y axis (running time, second). Observed speed-ups are indicated for individual analysis stages and overall running time for end-to-end pipeline on IGAP GWAS data (Lambert *et al.*, 2013).

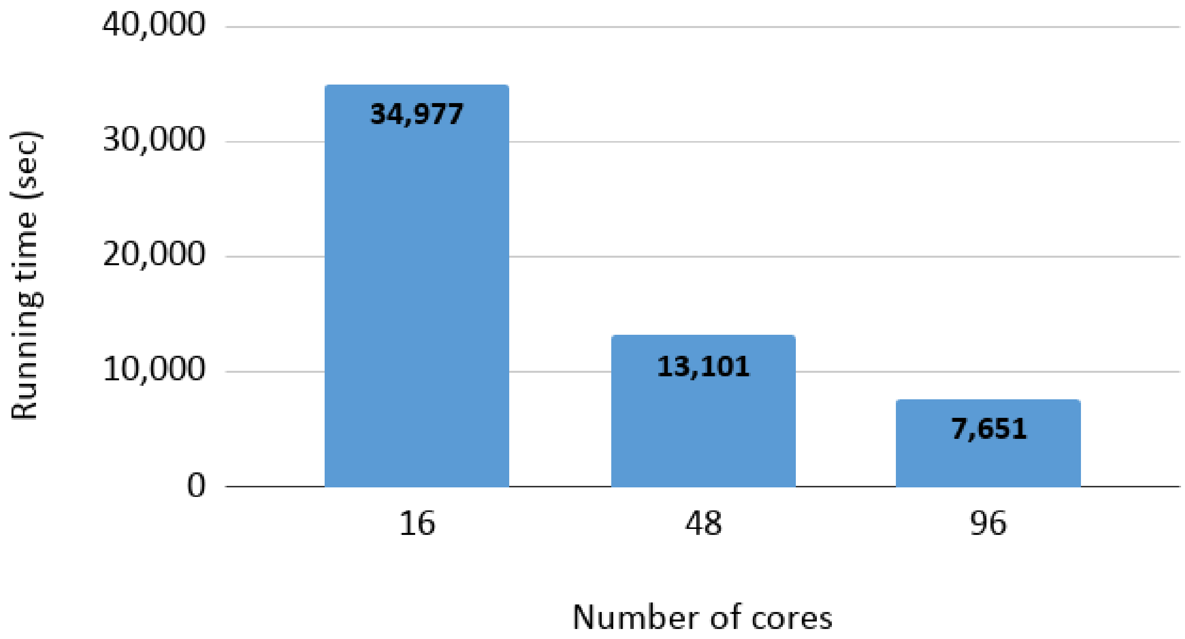


**Figure S3.** Scalability of SparkINFERNO pipeline. SparkINFERNO pipeline scales linearly with the number of computing cores. IBD GWAS dataset (Liu *et al.*, 2015) with 11,555,676 SNPs (10,742 genome-wide significant SNPs with $p<5\times10^{-8}$; 886 independent loci) was used to evaluate SparkINFERNO running time. 15,343 candidate IBD-associated SNPs were overlapped with 1.8 Billion intervals (genomic features) at the rate of 18-30 Billion intervals/second for 16, 48 and 96 cores. 291,015 co-localization tests across 44 tissues and 886 loci at the rate of 8, 23, and 40 tests per second for 16, 48 and 96 cores.

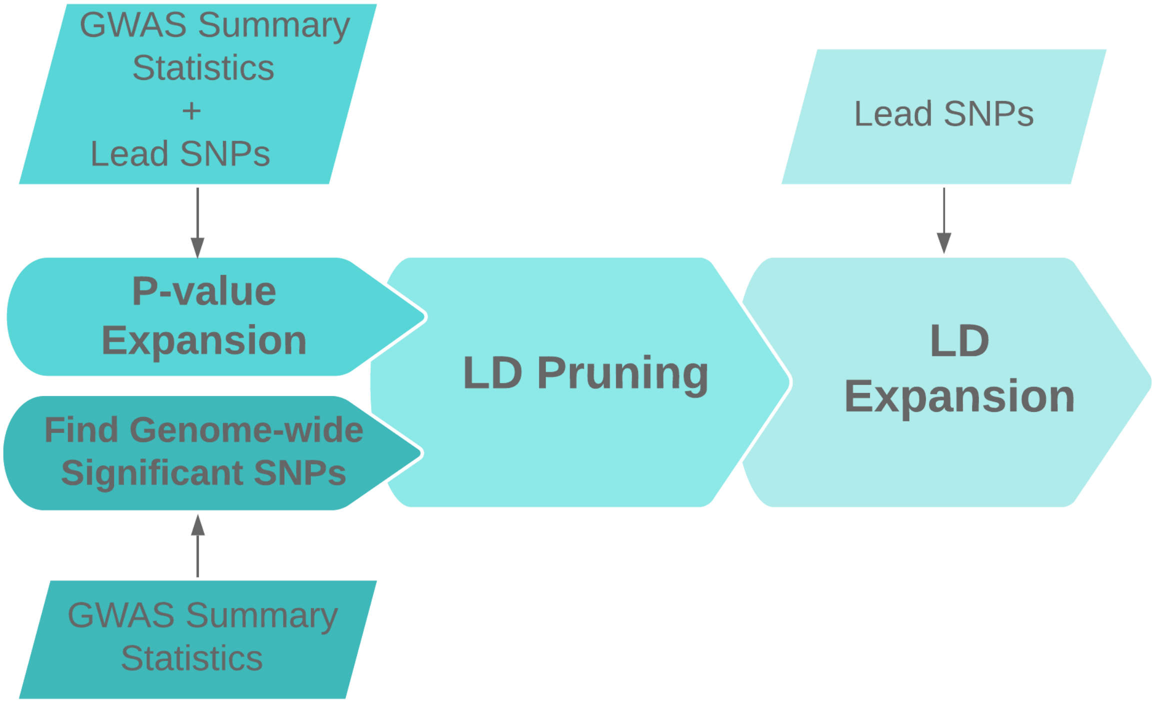


**Figure S4**. Pre-processing and candidate set generation. Lead SNPs (user-provided) and/or genome-wide significant variants are first LD pruned to identify independent signals, and then LD expanded to form a candidate set of potentially causal variants (see Pre-processing section under Supplementary Methods for details).

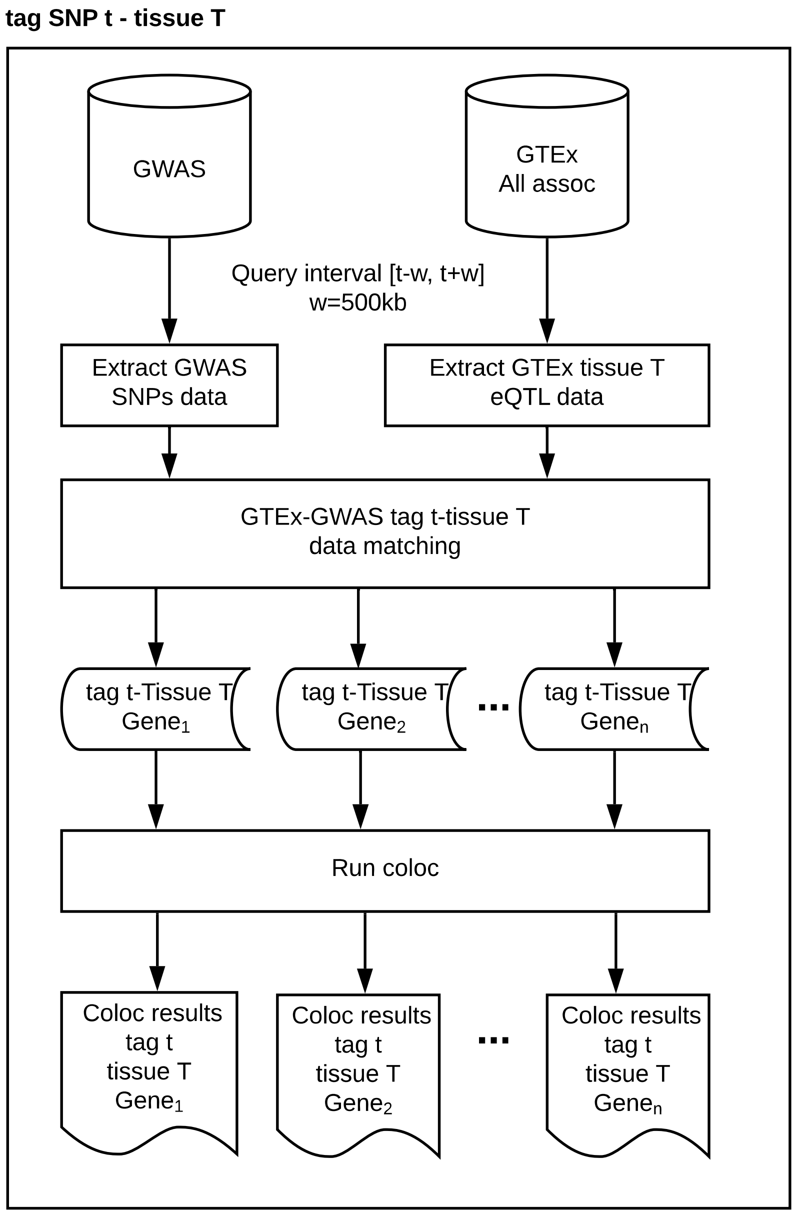


**Figure S5.** Spark-INFERNO scalable co-localization analysis architecture. Co-localization analysis (eQTL + GWAS) is parallelized across tag SNPs, tissues, and candidate genes. Shown is the co-localization analysis for a single tag variant *t* in a tissue *T*. All GTEx eQTL and GWAS signals (SNPs) are first extracted, matched, and then further split by gene to produce a set of tag t/tissue T/gene g datasets for co-localization analysis. Each co-localization test for tag t/tissue T/gene g uses R coloc package to produce per locus co-localization summary and detailed per-SNP co-localization results.

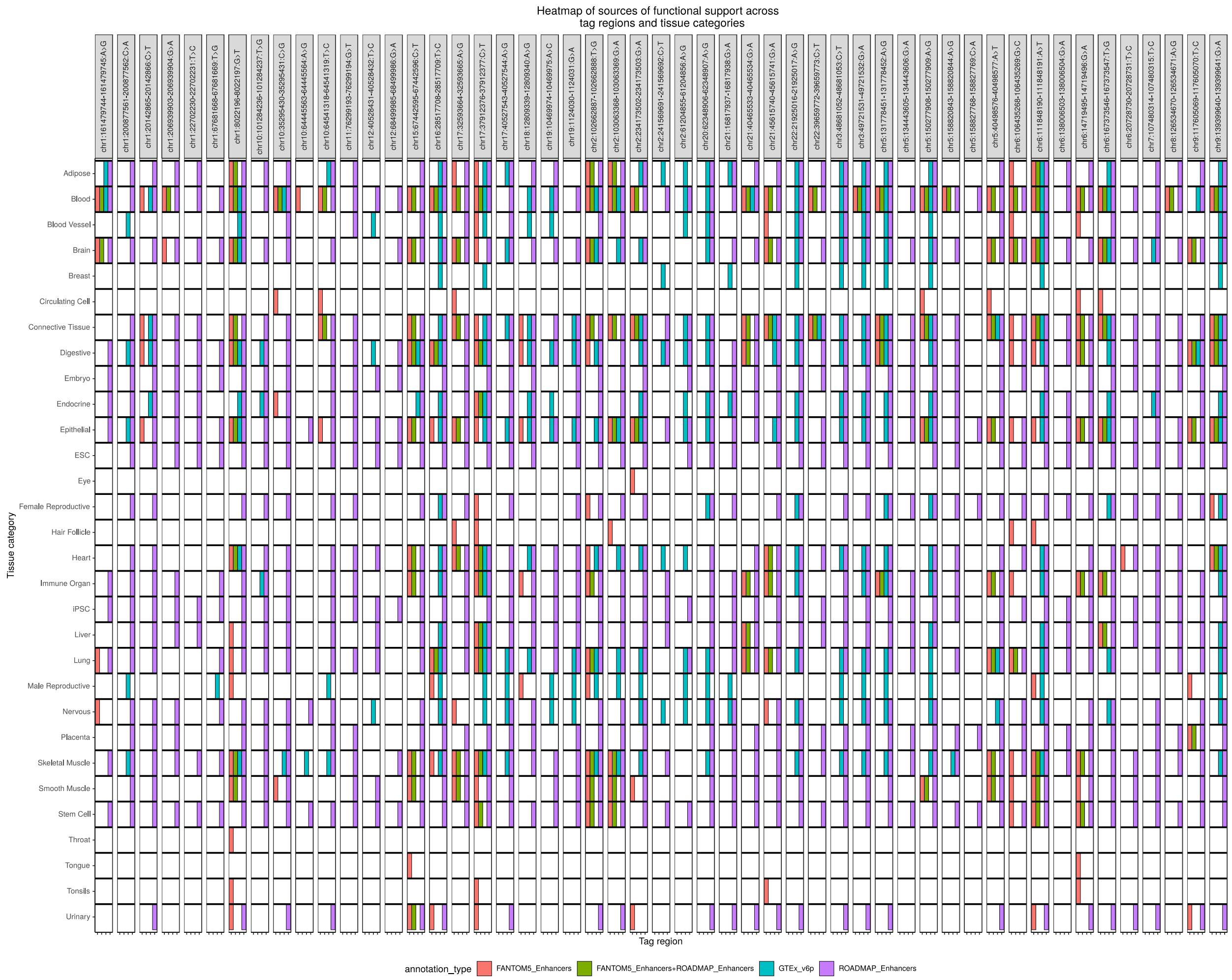


**Figure S6**. Summary of tissue category and functional genomics overlaps across tag regions (IBD GWAS).

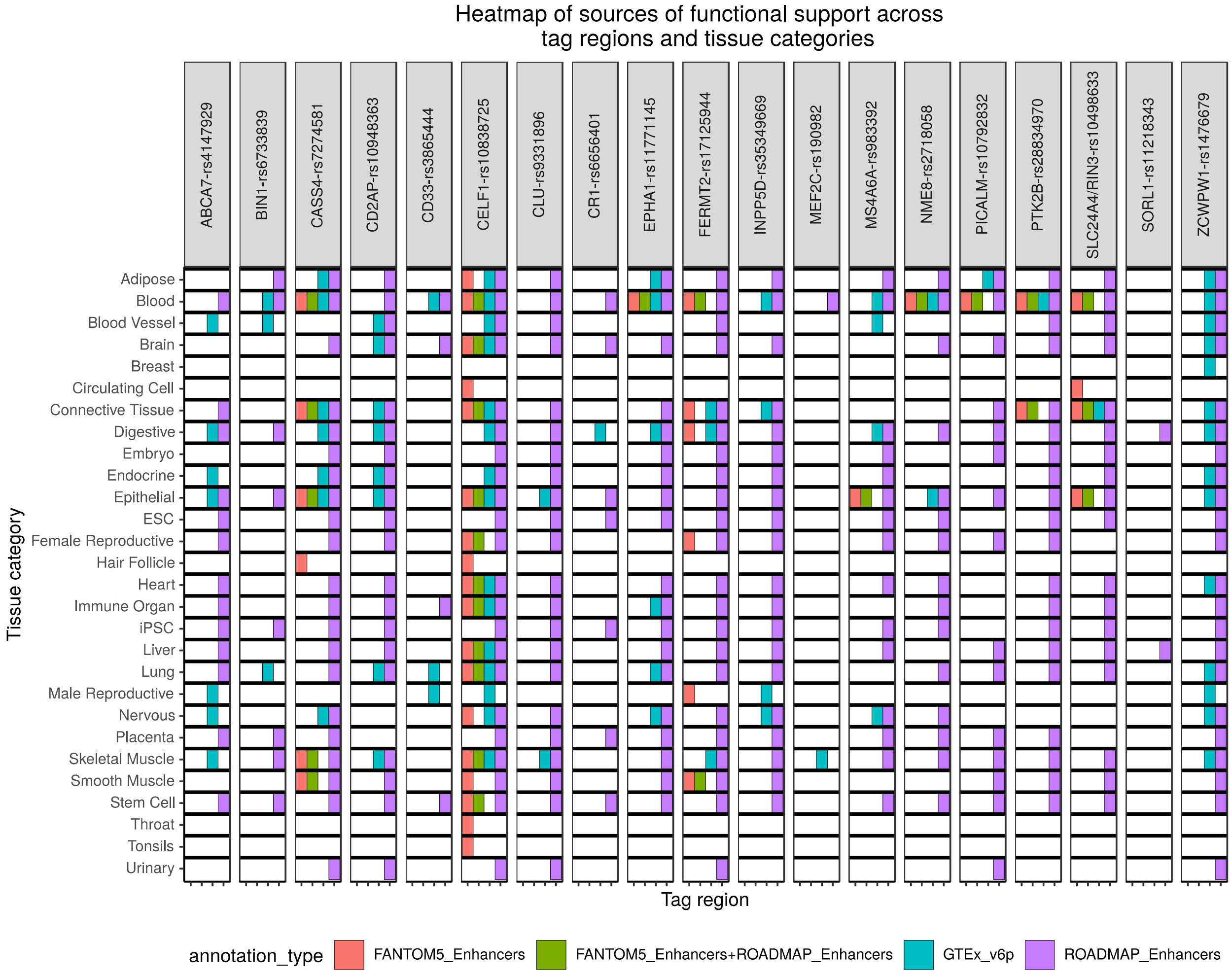


**Figure S7**. Summary of tissue category and functional genomics overlaps across tag regions (IGAP AD GWAS).

### Supplementary Methods

#### Annotation resources

Annotations for mRNA genes and repeat elements are obtained from UCSC Genome Browser (Tyner *et al.*, 2017) (knownGene, kgXref, RMSK tables).
The annotation information for long non-coding RNAs is based on LNCipedia v4.1 (Volders *et al.*, 2015); information on tissue-specific small RNA loci is based on DASHR (GRCh19/hg19, GRCh38/hg38 genomes) (Kuksa *et al.*, 2018; Leung *et al.*, 2016);
Tissue-specific enhancer loci were obtained from FANTOM5 (Andersson *et al.*, 2014). Tissue-specific chromatin state information is obtained from Roadmap (Consortium *et al.*, 2015). Transcription factor binding information was obtained from Homer project (Heinz *et al.*, 2010). Expression QTL datasets were obtained from GTEx project (Aguet *et al.*, 2017).

#### SparkINFERNO pipeline

The SparkINFERNO pipeline involves the following steps:

1. Confirming the correctness of SNPs from GWAS summary statistics and resolving alleles
2. Identifying genome-wide significant GWAS signals (SNPs) from GWAS summary statistics if lead SNPs are not defined.
3. If the defined lead SNPs are provided, for each lead SNP, using P-values of nearby SNPs to expand additional SNPs from GWAS. Otherwise, using genome-wide significant GWAS signals can skip this step.
4. Building a candidate set of all potentially causal SNPs and loci using linkage disequilibrium (LD) pruning of top SNPs from the previous step followed by LD expansion.
5. Performing overlap of candidate SNPs from step 4 with functional genomics data collections.
6. Performing statistical ranking of the significance of overlaps using permutation/randomization test (enrichment analysis).
7. Conducting co-localization analysis of the GWAS + eQTL (GTEx) data.
8. Summarizing results for each input signal (SNP) by tissue, and annotation category for each of the data collections.
9. Meta-table construction, evidence concordance analysis: integrate / meta-analyze results across data collections and above genomic analyses.
10. Identifying co-localized, concordant signals and output prioritized SNPs, enhancers, tissues, target genes.

#### Pre-processing

Pre-processing procedure includes several data processing steps for the GWAS summary statistics. First, GWAS QC step is conducted to confirm the correctness of GWAS SNPs by using allele frequency and rsID information as well as applying 1000 Genomes Project with a chosen super population to resolve effect and non-effect alleles as corresponding reference and alternative alleles. Next, finding top SNP step is performed to identify genome-wide significant

GWAS signals. If lead SNPs are defined, P-value expansion step is applied to expand SNPs within 500kb both sides of each lead SNP with a P-value threshold. Then, identifying independent signals/loci by LD pruning, and constructing a set of potentially causal variants by LD expansion of independent signals/loci. Finally, output potentially causal variants as a standard BED file for a downstream processing of functional data overlap and co-localization analysis.

**Functional data overlap**

Overlap is performed by using Spark-based mapping of input data (GWAS) against each of the datasets in the SparkINFERNO functional genomics data repository. Each individual overlap operation is performed using Giggle. Overlaps are further joined with attributes (e.g., tissue type, tissue category, and total interval number, etc) of tracks in the data repository, and independently grouped by intervals, tissues, and tracks to summarize the overlaps. Resulting Spark overlap data frames are saved as a table in TSV format.

**Enrichment analysis**

To assess significance of overlaps with functional genomic annotations and datasets, SparkINFERNO computes empirical p-values using randomization test by running overlaps pre-specified number of times (default: 1000) using shuffled data.

**Co-localization analysis**

Tissue-specific co-localization analysis is computationally demanding with the complexity scaling as (Number of tag regions (LD blocks) * region size (Number of SNPs) * Number of eQTL Genes * number of tissues). SparkINFERNO implementation (Figure S5) fully exploits distributed, parallel nature of Spark architecture to achieve highly efficient, distributed co-localization analysis.

**Meta-table construction**

All interval summaries and the co-localization results are collapsed as a meta-table. The meta-table contains SNPs information from the input BED file, tissue types (and count), tissue categories (and count), concordant tissues, functionally genomic partitions, motif sequences and positions from each functional annotation, and co-localized signals from each eQTL gene and tissue, etc.

**SparkINFERNO input**

The required input for SparkINFERNO is either one of the following: 1) GWAS summary statistics + defined lead SNPs, 2) GWAS summary statistics only, or 3) defined lead SNPs only. The different input combination leads to different procedure in the pre-processing step. The other required input is annotation tracks/data repository (Figure S1).

**SparkINFERNO output**

After running an entire SparkINFERNO pipeline with the input combination of GWAS summary statistics + defined lead SNPs or GWAS summary statistics only, a meta-table named "coalesced_coloc_stats_metatable.tsv" is produced with results across all functional annotations and co-localization analysis. Briefly, the meta-table includes the input SNP information, tissue type (and count), tissue category (and count), concordant tissues, functionally genomic partitions, motif sequences and positions from each functional annotation, and co-localized signals from each eQTL gene and tissue. The detailed column information and description are shown in Table S3 (inferno_metatable_description.xlsx). Note that as GWAS summary statistics are required to conduct the co-localization analysis, when only lead SNPs are provided, SparkINFERNO generates the simplified meta-table called "coalesced_metatable.tsv" without co-localization result.

**SparkINFERNO evaluation**

We systematically evaluated SparkINFERNO using publicly available GWAS datasets (IBD and IGAP AD). To evaluate scalability, we tested SparkINFERNO running time as a function of number of computing cores (Figure S1) on Linux AWS server instances (m5.4xlarge – 16 cores / 64 GB memory, m5.12xlarge – 48 cores / 192 GB memory, and m5.24xlarge – 96 cores / 384 GB memory). We also analyzed overall and individual stage running time on IGAP data (Figure S2) in comparison with original INFERNO implementation (Amlie-Wolf *et al.*, 2018).
